# 3Cnet: Pathogenicity prediction of human variants using knowledge transfer with deep recurrent neural networks

**DOI:** 10.1101/2020.09.27.302927

**Authors:** Dhong-gun Won, Kyoungyeul Lee

**Author notes:** e-mail address: Dhong-gun Won, Kyoungyeul Lee.

## Abstract

Thanks to the improvement of Next Generation Sequencing (NGS), genome-based diagnosis for rare disease patients become possible. However, accurate interpretation of human variants requires massive amount of knowledge gathered from previous researches and clinical cases. Also, manual analysis for each variant in the genome of patients takes enormous time and effort of clinical experts and medical doctors. Therefore, to reduce the cost of diagnosis, various computational tools have been developed for the pathogenicity prediction of human variants. Nevertheless, there has been the circularity problem of conventional tools, which leads to the overlap of training data and eventually causes overfitting of algorithms. In this research, we developed a pathogenicity predictor, named as 3Cnet, using deep recurrent neural networks which analyzes the amino-acid context of a missense mutation. 3Cnet utilizes knowledge transfer of evolutionary conservation to train insufficient clinical data without overfitting. The performance comparison clearly shows that 3Cnet can find the true disease-causing variant from a large number of missense variants in the genome of a patient with higher sensitivity (recall = 13.9 %) compared to other prediction tools such as REVEL (recall = 7.5 %) or PrimateAI (recall = 6.4 %). Consequently, 3Cnet can improve the diagnostic rate for patients and discover novel pathogenic variants with high probability.

## Introduction

Missense variants refer to variants caused by missense mutations in which an amino acid of a protein residue is replaced by another amino acid due to the nucleotide change of a gene. Missense mutations are common which correspond to 83% of non-synonymous mutations in the population and the pathogenicity is less severe than nonsense variants or frameshift variants in most cases ^1^. Nevertheless, many rare diseases are caused by the missense mutations such as developmental disorder, heart malformation, and many kinds of syndromic disorders ^2–4^. Recent improvements in the Next Generation Sequencing (NGS) enable finding a massive number of mutations in patients. Given the frequent occurrence of missense variants, it is not surprising that missense variants are commonly found in the genome of patients suffering rare diseases ^5^. However, clinical pathogenicity of a missense variant is difficult to predict, and it is necessary to comprehensively consider the effect of the variant on the protein, on the cell, and eventually on the human body ^6^. Also, identifying the true disease-causing variant among many missense variants is crucial for the diagnosis of patients. Therefore, analyzing the effect of missense mutations is an important and challenging problem for clinical uses.

There is a standard guideline for diagnosing patients based on the interpretation of sequence variants recommended by the American College of Medical Genetics and Genomics, that is, the ACMG guideline ^7^. Based on the guideline, pathogenicity of each variant from patients has been reported to a public database, ClinVar ^8^, as one of the five classes including pathogenic (P), likely pathogenic (LP), uncertain significance (VUS), likely benign (LB), and benign (B). However, In ClinVar, the number of missense variants recorded with known pathogenicity and reliable confidence is less than 100,000. As the possible number of missense mutations within the human genome is 82,755,468 according to dbNSFP ^9^, the pathogenicity of missense mutations are rarely known. Also, it takes a lot of time and effort until the disease-causing variant is confirmed and the patient is diagnosed, which leads to high failure rate of diagnosis and delayed treatment for patients ^10^. As mentioned, due to the importance of missense mutations in rare diseases, there has been unmet needs to specify the pathogenicity of VUS mutations found in the patient genome ^11^. If a prediction algorithm can predict the disease-causing variants in advance, it can reduce the time and cost required for diagnosis considerably. PP3 is one of the standards in the ACMG guideline which means in-silico assessment criterion. The importance of PP3 is continuously growing because the assessment of missense variants depends largely on in-silico prediction ^11^.

In the meantime, with the development of machine learning, algorithms and services based on artificial intelligence have been developed in many fields. In the field of computational genomics, various attempts have been made to develop artificial intelligence-based diagnostics using rapidly increasing genomic data. Some of the attempts have been made to predict the pathogenicity of variants using machine learning algorithms. Representatively, REVEL developed a random forest algorithm that incorporates various pathogenicity predictors to build an integrated predictor for missense variants. CADD ^12^ is another ensemble method which uses linear regression to integrate different scoring tools. FATHMM makes use of evolutionary information to recognize evolutionarily well-conserved variants ^13^. VEST4 ^14^, POLYPHEN2 ^15^, and SIFT ^16^ are other well known prediction tools to predict the functionality change based on Random Forest, Naive Bayes, and statistical method, respectively. DANN is the first artificial neural network which uses 949 features to predict the pathogenicity of nonsynonymous variants ^17^. However, there have been critical issues about circularity of the conventional predictors caused by making use of scores of other tools ^18^. Circularity can lead to an overlap between training and evaluation datasets and consequently result in overfitting. Also, the shortage and bias of clinical data triggers overfitting of AI machines to previous knowledge.

On the one hand, there was a novel attempt to solve this problem by applying sequence based pathogenicity prediction. PrimateAI makes use of sequences of wild-type and mutant proteins to compare the difference and decide the pathogenicity of mutations using Convolution Neural Network (CNN) ^19^. Such an approach which utilizes the protein sequences for pathogenicity prediction is promising because it can avoid the circularity problem and overfitting to previous knowledge. However, Compared to the adequate number of data to train deep neural networks effectively, the number of clinical data available in ClinVar is relatively small. Instead, PrimateAI made use of common human variants and variants from primates as benign data while simulated variants based on trinucleotide context were used as unlabeled data.

The previous studies using deep learning had some limitations for clinical use of the predictors. First of all, the performance of pathogenicity prediction for clinical data was too low compared to random forest models such as REVEL or VEST. This may be because deep neural networks require massive amounts of data for effective training compared to random forests. When a deep neural network is solely trained by clinical data from ClinVar, the model would be overfitted to the data so that it can hardly interpret novel pathogenic variants. Nevertheless, the usage of clinical data from patients diagnosed based on the ACMG guideline is essential to build accurate pathogenicity predictors. Deep neural networks trained without clinical data show low accuracy to discriminate pathogenic variants from benign variants for human mutations. We overcame such limitations of deep neural networks by transferring the knowledge obtained from evolutionary conservation of protein sequences with Recurrent Neural Networks (RNNs).

Knowledge transfer in deep learning indicates utilizing data of similar tasks to train a specific task more effectively, which generally has insufficient number of data ^20^. In this study, we utilized knowledge transfer to integrate three different types of data: clinical data, common variants, and conservation data. Clinical data consist of pathogenic variants and benign variants reported to ClinVar based on the ACMG guideline. Common variants indicate variants frequently observed in the general human population which can help to find benign variants of patients ^21^. Conservation data is referred to as the simulated variants we generated based on the trinucleotide context and evolutionary conservation. We compared three different strategies for knowledge transfer, which are single-task learning, multi-task learning, and transfer learning, to find the best practice for training. The optimized predictor, named as 3Cnet, was able to evaluate the impact of missense variants much more accurately than other deep neural networks (PR-AUC = 0.885 vs. 0.791) and better find the disease-causing variants of rare disease patients compared to ensemble models (recall = 0.139 vs. 0.075). 3Cnet is the first RNN based pathogenicity predictor which learns the effect of variants in the context of protein sequences. Additionally, we applied knowledge transfer of conservation data for other challenging problems regarding variant pathogenicity such as protein stability change and gene expression change in the cell. We found that conservation data can also improve the prediction accuracy for those problems by reducing overfitting of given data.

## Methods

### Curatation of clinical data from ClinVar database

We performed 5 fold cross validation for 65,239 missense mutations in ClinVar database (released in April, 2020). We first curated the mutations in ClinVar to find missense mutations with known molecular consequences and having reliable review status. For that purpose, we collected mutations of which molecular consequence was ‘missense variant’ and excluded mutations having unreliable review status such as ‘no assertion for the individual variant’, ‘no assertion criteria provided’, and ‘no assertion provided’. As our prediction algorithm makes use of protein sequences around mutation sites as input features, each variant was represented as the HGVS term in which the transcript ID and the mutation information were given ^22^. For the transcript ID, the canonical transcript in Refseq database ^23^ was referred. Then, each missense mutation was transformed into data representing a protein sequence composed of 201 amino-acids centered around the mutation site. Sequence data for both wild-type protein and mutant protein were generated to compare the difference in the context of amino-acid sequences.

There could be multiple reports for a single variant and sometimes they could be in conflict (ex. one report is pathogenic while the other report is benign). Therefore, the pathogenicity of each variant was determined by integrating pathogenicity reports from ClinVar for the same variant. There are 5 labels for pathogenicity which are pathogenic (P), likely pathogenic (LP), variants with uncertain significance (VUS), likely benign (LB), and benign (B). We set a standard to define pathogenicity for each variant from multiple reports. When there are any reports saying a variant is pathogenic or likely pathogenic, we consider the variant is pathogenic except for the cases in which there are contrary reports. Similarly, a variant with any reports saying the variant is benign or likely benign is considered as a benign variant. We removed variants having contrary reports and variants of which reports are VUS only. As a result, we got 22,278 pathogenic variants and 48,580 benign variants from the ClinVar database. The input features of the clinical data are sequence representations of wild-type protein and mutant protein, while the output feature is a binary label for pathogenicity.

### Augmentation of clinical data using common variants in GnomAD

We used the GnomAD database to gather missense mutations frequently observed in the general population, namely common variants ^24^. Such variants are found in the genome of a number of general people which means they are probably benign variants. Common variants were used as benign variants to train predictors to get better precision by reducing false positives ^25^. Firstly, Common variants having allele frequency (AF) higher than 0.1 % were collected. Then, those variants were represented as the HGVS term based on the canonical transcripts as we did for ClinVar database. Only variants not included in ClinVar database were curated as common variants to avoid conflicts and overfitting due to duplicate samples. In total, 59,375 variants were found to be common variants and their pathogenicity were labeled as benign. As the benign variants including data from ClinVar were much bigger than pathogenic variants (22,278 vs. 107,955), pathogenic samples were augmented 4 fold during training for balance. After removing a few transcripts inconsistent with the reference sequence of HG19 ^26^, 18,942 unique transcripts were found in the curated data in total.

### Generation of conservation data using Multiple Sequence Alignment

For 18,942 transcripts included in the data we curated, multiple sequence alignment (MSA) was constructed to see the evolutionary conservation patterns of those proteins. For each transcript, the sequence was transformed into FASTA format. Then, a hidden markov model based algorithm, HHblits ^27^, built MSA from the query sequence using UniRef30 (version 2020.02) as a sequence database ^28^. Those MSA results were utilized for two purposes: to derive input features of the networks and to generate simulated variants reflecting evolutionary constraint. Among the sequences aligned with the query sequence, only sequences having over 30% identity and 80% overlap with the query remained. In addition, only residues aligned with more than 10 sequences except for gaps were utilized for the following processes.

The number of variants from clinical data is 130,233 even if common variants are included as benign samples. The number is not enough for training a deep neural network having more than 10,000 features as sequence inputs. Therefore, to avoid overfitting that could occur when the amount of data is not sufficient, more mutation data were needed to train our model without bias. Based on the conservation patterns derived from MSA, we simulated variants which are frequently or hardly found in nature. First, we randomly created variants at each residue of the transcripts considering trinucleotide context in their genomes ^19^. Among those variants, we defined the variants that had never been found at the residue as pathogenic variants. On the other hand, the variants frequently found with the ratio higher than 50 % out of aligned amino acids were defined as benign variants. We randomly selected 10 % variants from all possible simulated variants to take reasonable cost for training. We named those variants as conservation data and utilized it to train models along with clinical data. Resulting numbers of mutations for each type of data are summarized in the table 1.

**Table 1.**
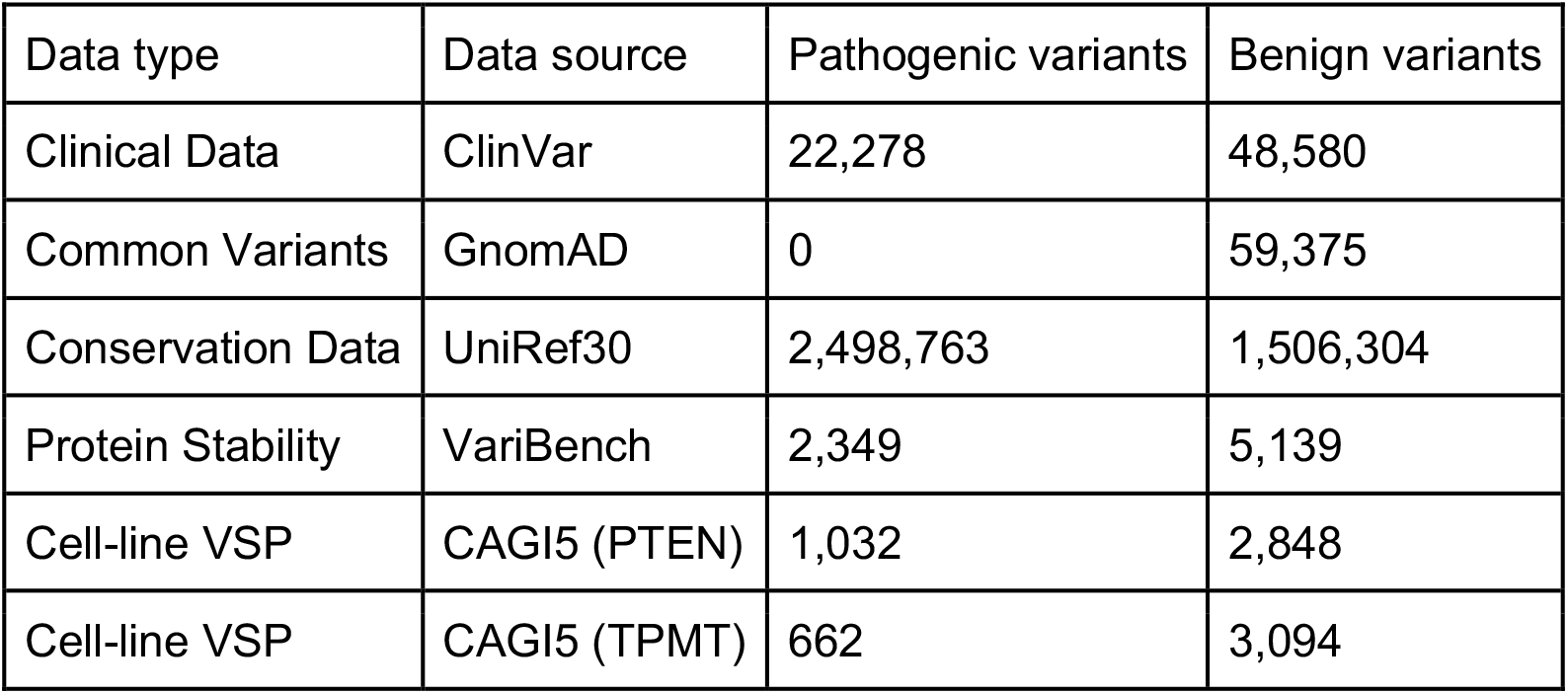
The number of pathogenic and benign variants for each type of data

### Featurization of sequences and MSA

To train deep neural networks, each variant needed to be transformed into the real value features which represent the amino-acid sequence around the mutation site. A sequence can be interpreted as sequential inputs of amino acids. Each type of amino-acid could have unique traits represented as a feature vector. We applied feature embedding to represent an amino acid in the form of a feature vector ^29^. As a result, a sequence was transformed into a feature matrix filled with sequential feature vectors representing amino-acids (Figure 1). As the length of overall sequences are various depending on the transcripts, we made use of only 201 residues around the mutation site (former 100 AAs to latter 100 AAs) to build the data. If the mutation site was too close to the start point or endpoint of the sequence so that some positions in the data cannot be matched to specific amino acids, such positions were filled with vectors with zeros (zero padding). Both wild-type sequence and mutant sequence were transformed into feature matrices and used as input features.

**Figure 1.**
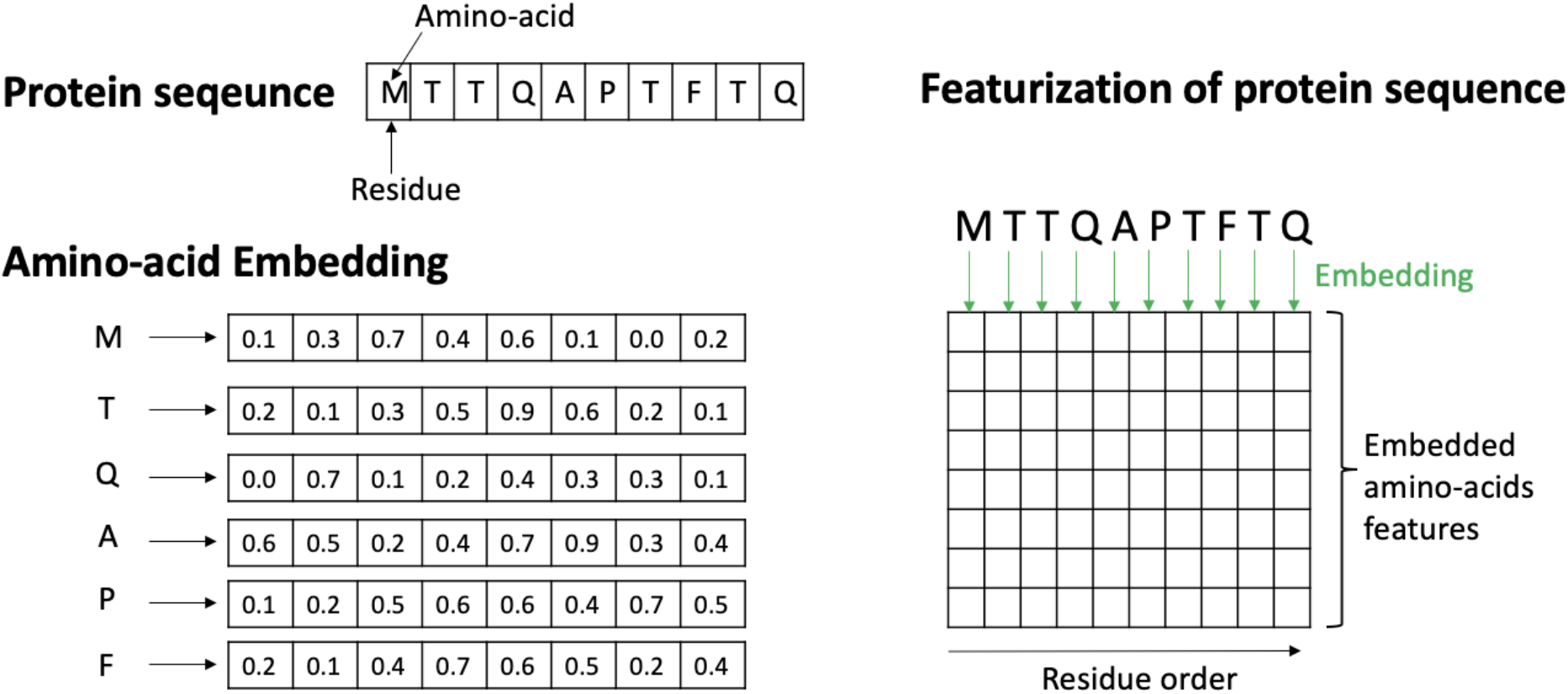
Transformation of a protein sequence into a feature matrix

Along with sequence data of the transcripts, we made use of conservation patterns in MSA as input features as well. As such features can represent evolutionary constraints imposed on the residues of transcripts, those features could be crucial for pathogenicity prediction. Therefore, each residue in the sequence was imposed with a vector containing the ratio of amino acids evolutionarily found in the residue based on MSA (figure 2). For those residues of which the number of aligned sequences is less than 10, values of the vector were filled with zeros. As a result, 8,726,858 residues from the 18,942 transcripts were imposed with ratio vectors while 3,693,654 residues were padded with zeros. As we did for sequence data, we selected 201 residues around the mutation site and built a feature matrix for those residues. The feature matrix is also used as input features to train pathogenicity predictors.

**Figure 2.**
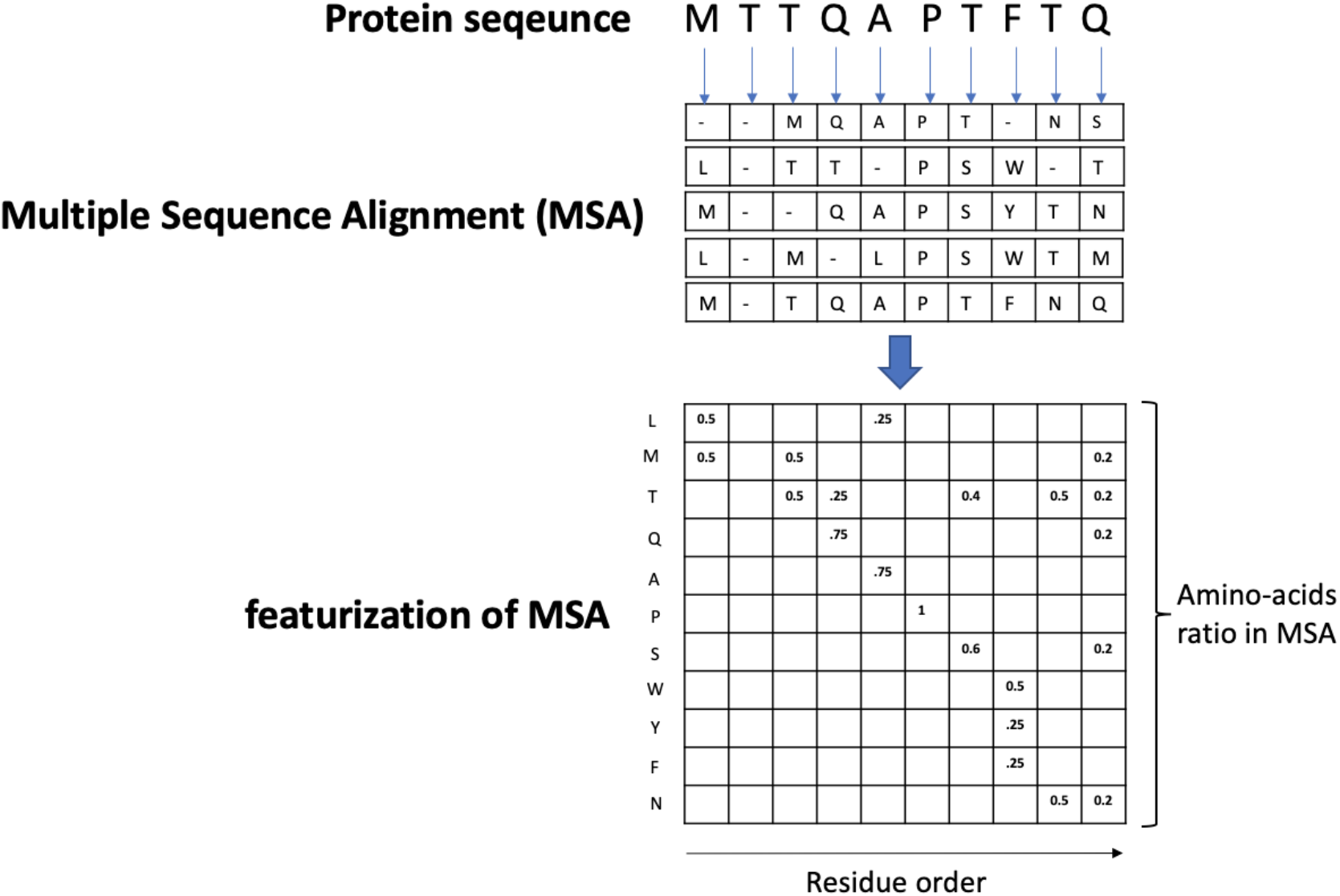
Transformation of MSA into a feature matrix

### Network architecture of the pathogenicity predictor

The model network we built can be divided into two modules based on their functions. The former part of the network is named as feature extractor, as the module extracts a feature vector of a variant based on the feature matrices of sequences and MSA (Figure 3). The latter part of the network is pathogenicity classifier which decides whether the variant is pathogenic or benign based on the feature (Figure 4).

**Figure 3.**
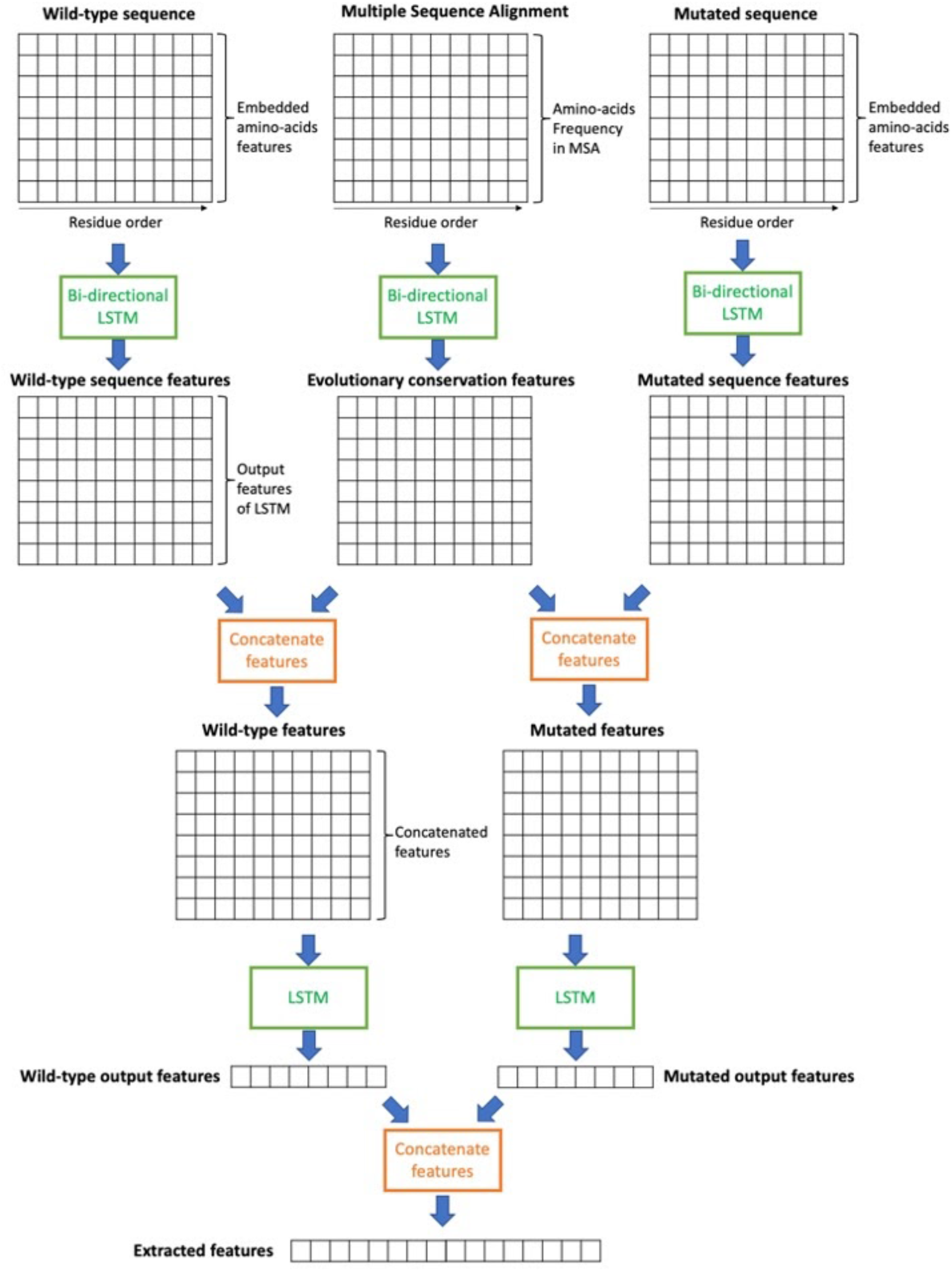
Deep recurrent neural network for feature extractor

**Figure 4.**
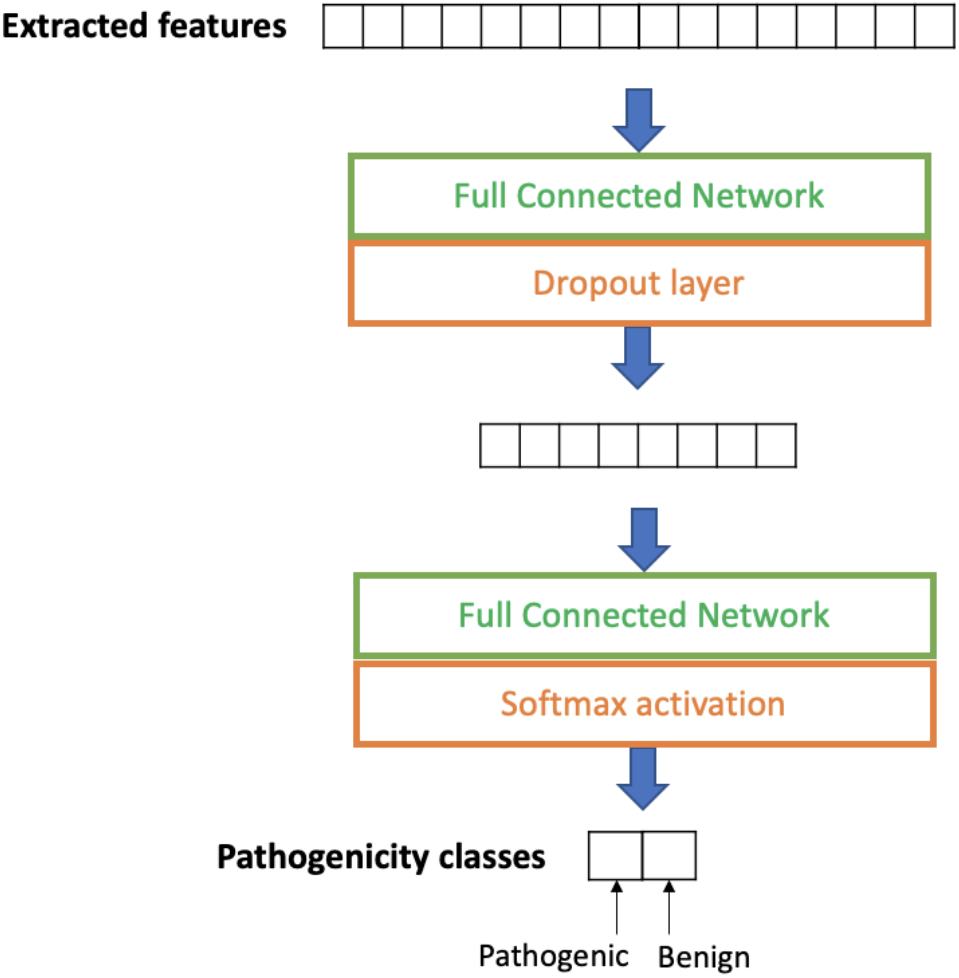
Neural network for pathogenicity classifier

The feature extractor is composed of 2 parallel layers utilizing Long Short-Term Memory (LSTM) networks, a type of RNN ^30^. The first layer consists of bi-directional LSTM networks which featurize three different input feature matrices independently, which are the wild-type sequence feature matrix, MSA feature matrix, and mutant sequence feature matrix. The output feature matrices can conceive the context of the sequences through the recurrent networks to consider the influence of one amino-acid to other amino-acids. Then, the output matrix from wild-type sequence and that of MSA are merged to make a concatenated feature matrix. Features for the same residue are concatenated so that the network can compare the amino-acid of the sequence with evolutionarily conserved amino-acids at that residue. output matrices of mutant sequence and MSA are also merged. Then, those two concatenated features (wild-type and mutant) are featurized once more using LSTM networks. In this time, however, only the last feature vector for each recurrent network remains. Finally, the output feature vector of wild-type and that of mutant are concatenated to become an extracted feature vector.

On the other hand, the pathogenicity classifier is composed of two fully connected (FC) layers. The first FC layer is expected to extract the difference between wild-type and mutant as a feature vector. a dropout layer was applied to the first layer to avoid overfitting. Then the final FC layer is used to decide pathogenicity of a variant by applying softmax activation classifying the variant into pathogenic class or benign class. The binary cross entropy between labels and the predicted classes becomes the loss function of the network.

### Training models using various data and knowledge transfer

At first, we trained the model only using clinical data which were curated from ClinVar. Also, we trained the model with the clinical data along with common variants gathered from the GnomAD database which were used as benign samples. We checked the performance of the models using 5 fold cross validation of the clinical data. Only clinical data from ClinVar were used as the test set of cross validation. In another model, conservation data generated from MSA were used to train a predictor. As the patterns of evolutionary conservation are widely used to predict pathogenicity of genomic mutations, the model trained only with the conservation data might be able to predict the pathogenicity to some extent ^12,13,15^. Note that, in this study, hyperparameters of the network were not optimized to the data to compare different architectures and test data without bias. Also, we tried to transfer the knowledge obtained from evolutionary conservation to train clinical data without overfitting. Three different learning methods were adapted for knowledge transfer, which were single-task learning, multi-task learning ^31^ and transfer learning ^32^ (Figure 5).

**Figure 5.**
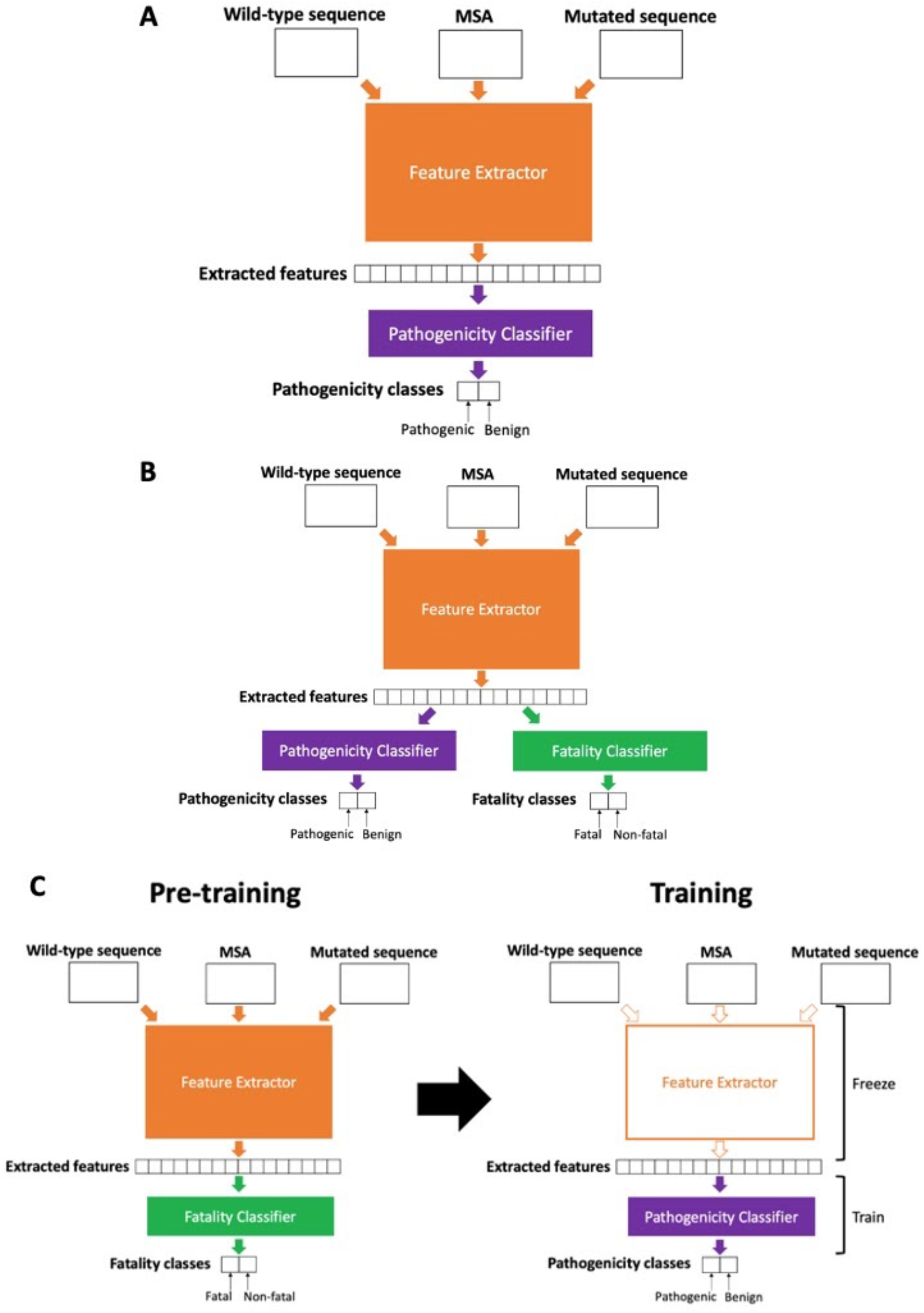
pathogenicity predictors using knowledge transfer (A) single-task (B) multi-task (C) transfer learning

For single-task learning, the conservation data and clinical data are trained alternately for a single network. As the number of the conservation data is much bigger than clinical data, clinical data are trained repeatedly to the network until all the conservation data are trained. On the other hand, for multi-task learning, the model utilizes the same feature extraction network, but independent pathogenicity classifiers are used for conservation data and clinical data. The extracted features are common features explaining both types of data, but the pathogenicity that each classifier predicts is separated. We named the classifier trained by the conservation data as fatality classifier because the pathogenic variants of conservation data are fatal which cannot be observed in nature. Finally, Transfer learning is a learning technique in which the model is pre-trained with the conservation data and then further trained with clinical data. By freezing layers for the feature extractor and only training the pathogenicity classifier, we can make use of features extracted based on conservation patterns.

## Result

### Cross validation for ClinVar variants

At first, we tested the performance of pathogenicity predictors using internal ClinVar variants using 5 fold cross validation (5FCV). Receiver operating characteristic (ROC) curve and precision-recall (PR) curve were measured and the area under curve (AUC) was calculated for each. Various types of training data and knowledge transfer were examined through cross validation and those AUC values were used as standards to compare different training strategies. Based on ROC-AUC and PR-AUC for various strategies, we chose the optimized prediction model for discovering pathogenicity of mutations. Note that only mutations in ClinVar data were cross-validated because only ClinVar mutations were labeled based on the ACMG guideline.

The Table 2 compares the performance of various training strategies in terms of ROC-AUC and PR-AUC. The result shows the predictor trained by clinical data along with common variants better predicts pathogenicity compared to the model trained only with clinical data. This result indicates that common variants with high allele frequency in the general population can be considered as benign mutations for training. Also, conservation data can be used to train a model as well, which was less accurate for prediction of clinical pathogenicity. Also, we checked the performance of the model trained by both clinical data and conservation data using various knowledge transfer techniques (see Method). In the case of multi-task learning, the prediction scores for validation were the scores from the layer trained by clinical data rather than conservation data.

**Table 2.** Accuracy, ROC-AUC, and PR-AUC of 5-fold cross validation for ClinVar data

| Model (CV5) | Accuracy | ROC-AUC | PR-AUC |
| --- | --- | --- | --- |
| Clinical data | 0.764 | 0.829 | 0.698 |
| Clinical data with common variants | 0.796 | 0.858 | 0.754 |
| Conservation data | 0.685 | 0.735 | 0.611 |
| Single-task learning | 0.761 | 0.878 | 0.803 |
| Multi-task learning | **0.832** | **0.893** | **0.824** |
| Transfer learning | 0.753 | 0.775 | 0.643 |

The result clearly shows that the performance of the model using multi-task learning between clinical data and conservation data is much better than other models using clinical data and conservation data respectively (Figure 6). We named the best model as 3Cnet. The superior performance of 3Cnet may come from the fact that the number of clinical data (65,239 mutations in ClinVar database) is insufficient to train a prediction model without bias ^33^. Although mutations from conservation data (about 40,000,000 variants) were artificially generated based on MSA and pathogenicity for each mutation had not been examined in clinical cases, such data could help to avoid overfitting of clinical data and enable the model to extract essential features reflecting evolutionary constraints of the proteins.

**Figure 6.**
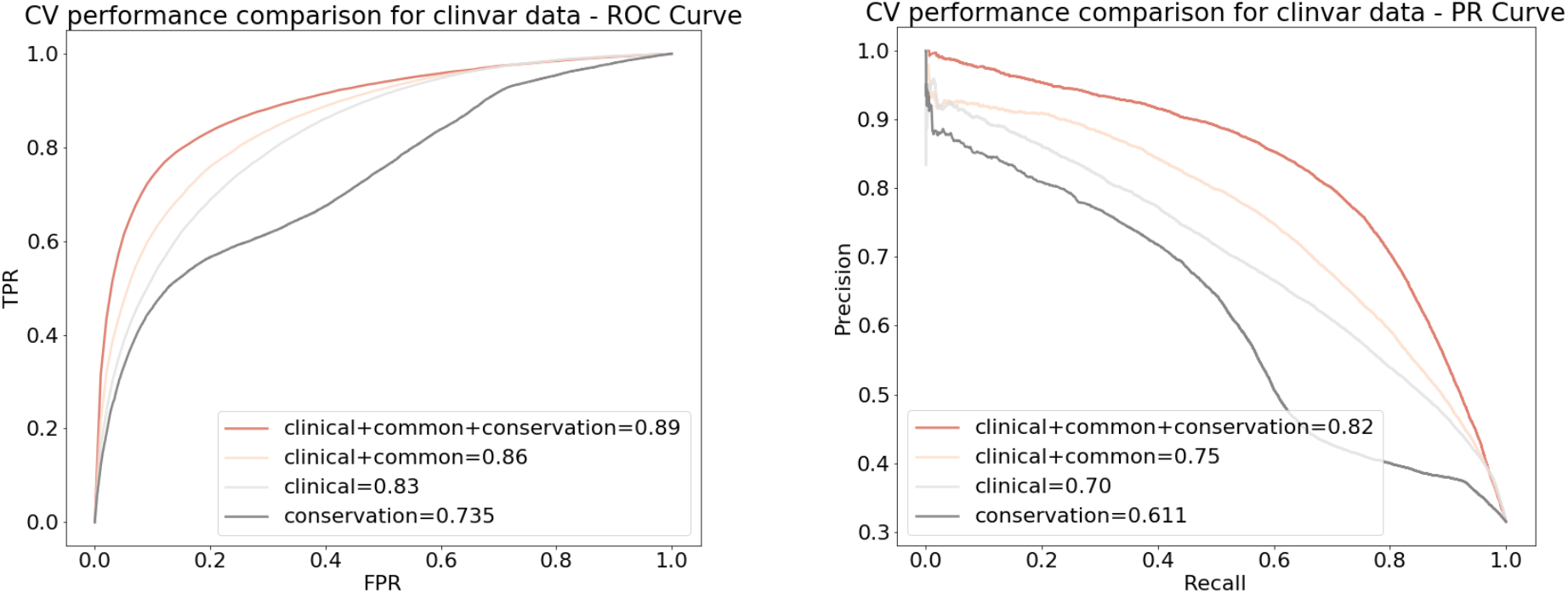
Cross validation performance for ClinVar variants. “clinical” means the model trained solely with ClinVar variants, while “clinical+common” indicates the model trained by clinical data along with common variants having AF > 0.1 % from GnomAD. “Conservation” is the model trained solely with conservation data. “clinical+common+conservation” stands for the model trained by multi-task learning between clinical data and conservation data. (left) ROC curve. (right) PR curve.

### External Validation for independent ClinVar variants

To compare other prediction tools with 3Cnet, we built an external dataset independent of the training dataset used for cross-validation ^34^. As the ClinVar dataset which we used as training data was released in April, 2020, we can make use of novel mutations in ClinVar released in August, 2020 as external data. After excluding all the overlapped mutations with training data, we gathered 10,262 pathogenic mutations and 12,041 benign mutations as external validation data. We tested the performance of 3Cnet using the external validation set and compared with other pathogenicity prediction algorithms. Scores for other tools including VEST4, PolyPhen2, SIFT, FATHMM, CADD, DANN, and REVEL are gathered from dbNSFP (version 4.0), while scores of PrimateAI are from Illumina website. Even though we were not able to specify every single mutation used to train those tools because of the use of commercially available database HGMD ^35^ and the circularity problem, we assumed that the external mutation data we built was distinctive from the mutations used to train other tools. The result in the figure 7 indicates that pathogenicity prediction of 3Cnet (PR-AUC = 0.885) is nearly as accurate as that of REVEL (PR-AUC = 0.912) or VEST4 (PR-AUC = 0.902) unlike other deep learning based predictors, PrimateAI (PR-AUC = 0.791) and DANN (PR-AUC = 0.649). Also, 3Cnet outperformed SIFT (PR-AUC = 0.841), PolyPhen (PR-AUC = 0.811), CADD (PR-AUC = 0.786), and FATHMM (PR-AUC = 0.783) which are widely used to confirm variant pathogenicity. The result indicates that 3Cnet can classify pathogenicity of variants more accurately compared to many other methods without making use of scores from other algorithms.

**Figure 7.**
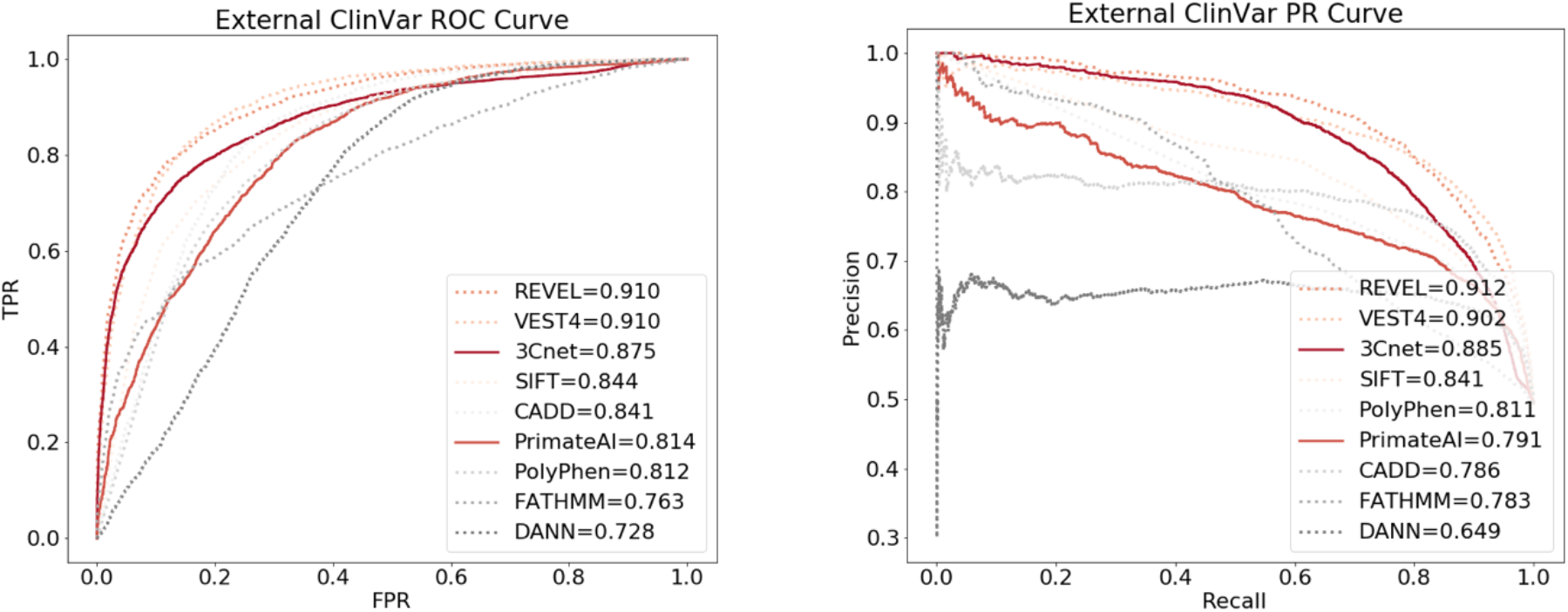
External validation performance for independent ClinVar variants and comparison with different prediction tools. Solid lines indicate the deep learning algorithms utilizing sequence inputs, 3Cnet and PrimateAI. Dotted lines mean the other conventional methods. (left) ROC curve. (right) PR curve.

### Patients data in diagnosis cases

We also checked how effectively 3Cnet can distinguish pathogenic variants from other missense variants of patients using the scores. As a matter of fact, such an application is more practical for diagnosis because we need to discover the disease-causing variant from a large number of missense mutations found in the genome of patients. We checked the cases of rare disease patients, and gathered missense variants found in the genome of 107 patients who were diagnosed based on the ACMG guideline ^36^. The disease-causing variants confirmed by medical doctors were annotated as positive samples while other missense variants from those patients were curated as negative samples. To test a reasonable number of variants, we removed variants having AF > 0.01 % in the general population. As some patients can have two disease-causing mutations for autosomal recessive diseases, there were 173 disease-causing variants and 29,473 uncertain variants to examine. We compared the PR curve of 3Cnet with those of REVEL and PrimateAI. ROC curves were not compared in this case because of the imbalance between positive and negative samples ^37^. Also, we compared the recall rate of the disease-causing variant for top-k variants of a patient, which represents the probability to find the true disease-causing mutations among high rank variants (Figure 8).

**Figure 8.**
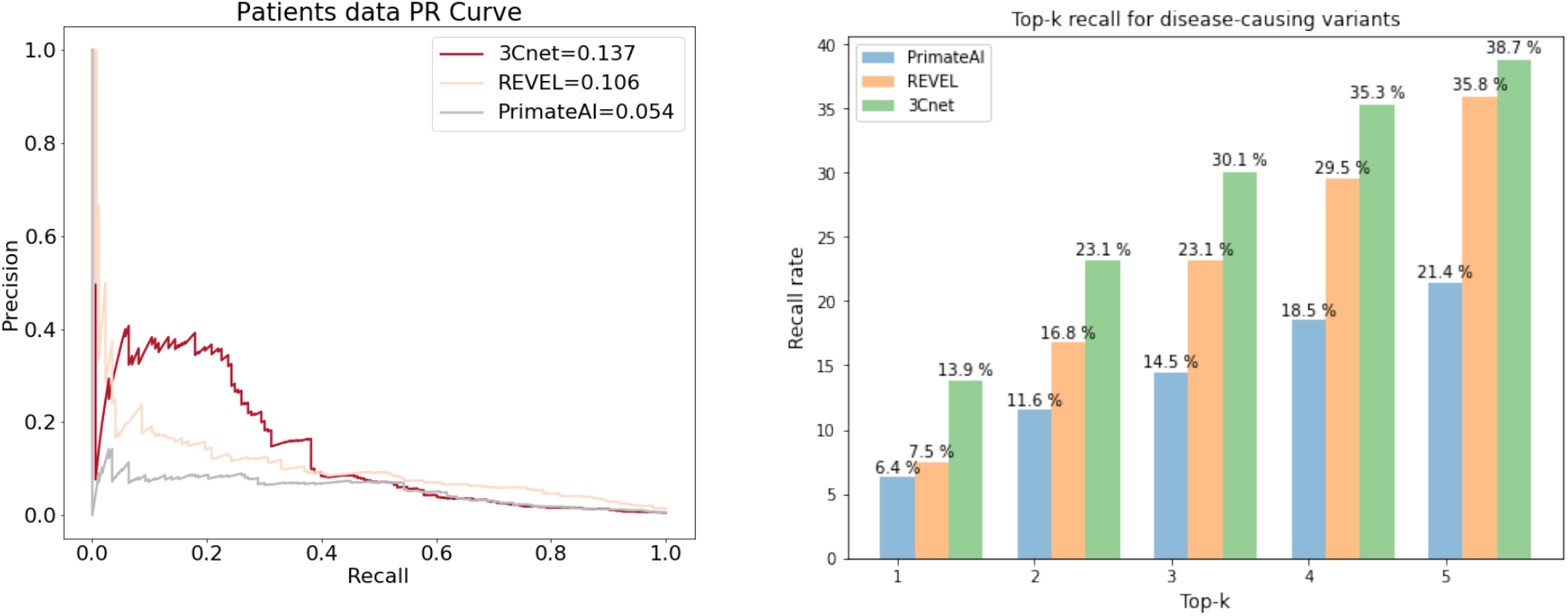
Comparison of performances for finding disease-causing variants among other missense variants of patients. (left) Precision-recall curve. (right) top-k recall

The result shows that 3Cnet can find disease-causing variants more effectively (PR-AUC = 0.137) compared to REVEL (PR-AUC = 0.106) or PrimateAI (PR-AUC = 0.054). 3Cnet shows 20 % recall rate maintaining a reasonable precision rate (around 40 %) compared to other methods (10∼15 %). However, the precision was high for REVEL when recall rate is less than 3%. Such a performance indicates that REVEL can verify pathogenic variants with high accuracy for a few variants but it neglects most of disease-causing mutations. In addition, top-k recall of 3Cnet was significantly higher than REVEL or PrimateAI representing the strength to find disease-causing variants of patients with better sensitivity.

### Prediction of protein stability change

We also tried to make use of the training strategy of 3Cnet to predict other types of pathogenicity such as protein stability change of mutations gathered from the VariBench database ^38^. The change in Gibbs free energy change (ddG) was used as the standard to measure protein stability change as it is a well known metric to predict the impact of a single point mutation ^39 40^. In general, ddG larger than 0.5 indicates destabilization of the mutated protein while ddG smaller than 0.5 is considered neutral or even more stable. In this study, we assumed that variants with ddG larger than 0.5 are pathogenic mutations while variants with ddG smaller than 0.5 are benign mutations. Consequently, there were 2,349 pathogenic variants and 5,139 benign variants in terms of protein stability.

Compared to the cross validation of clinical data described above, only clinical data is substituted into protein stability data and conservation data was utilized as same. Common variants were not used as benign samples in this practice. The performance of protein stability prediction was measured for different strategies using cross validation (Table 3 and Figure 9). The result shows that prediction performance was optimized when both protein stability data and conservation data were trained using multi-task learning. It indicates that conservation data we generated can be used not only for clinical data but for general purposes to train the impact of missense mutations. It also supports the thesis that conservation data can help training the pathogenicity predictor by reducing overfitting.

**Table 3.**
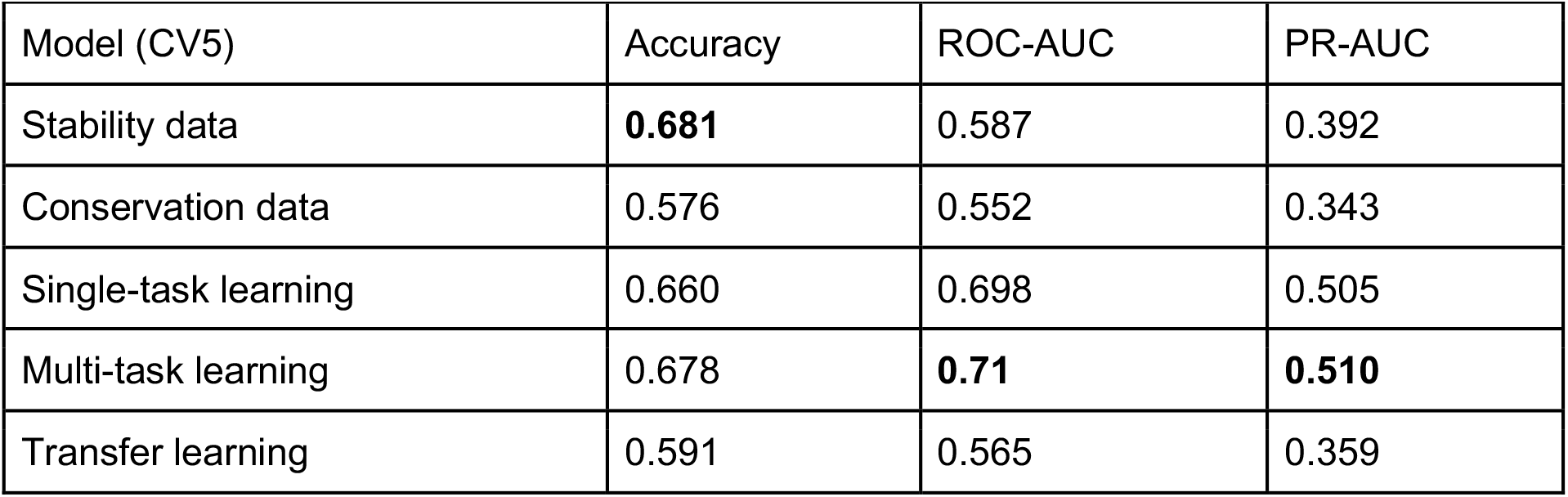
Accuracy, ROC-AUC, and PR-AUC of 5-fold cross validation for protein stability data

**Figure 9.**
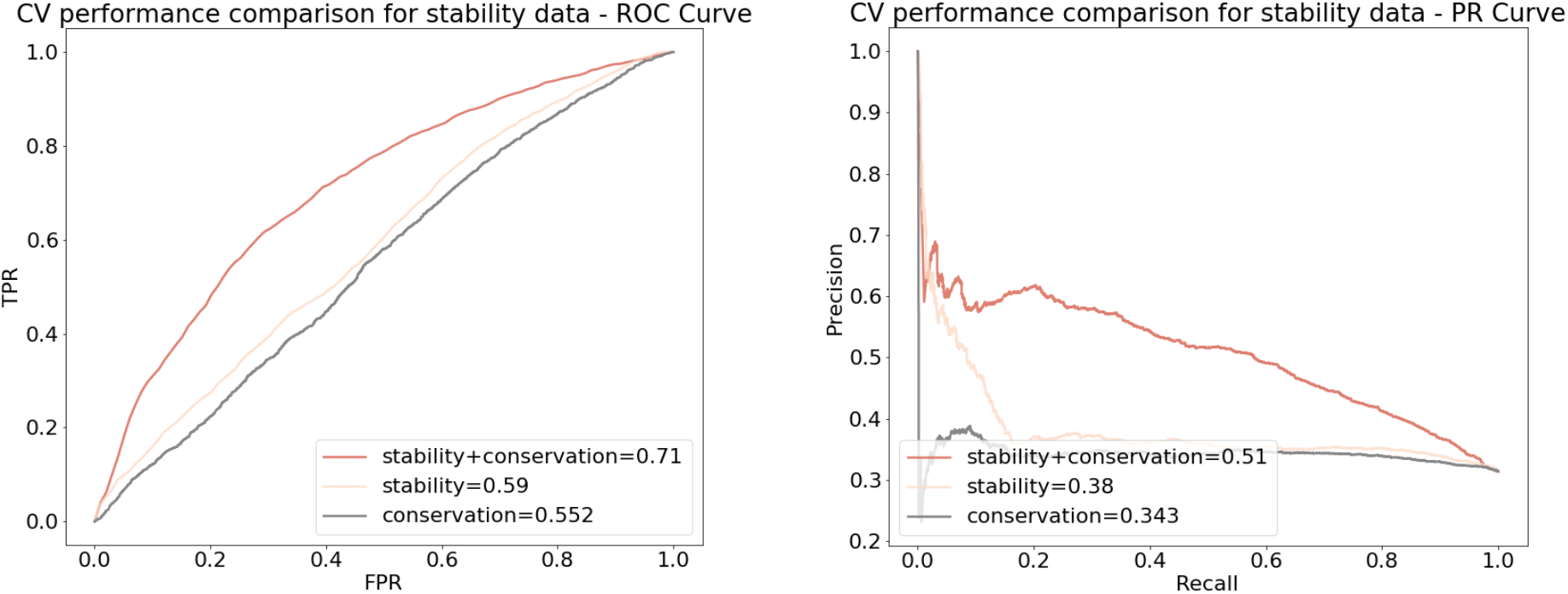
The performance of cross validation for protein stability prediction. “stability+conservation” means the model trained by multi-task learning between protein stability data and conservation data. (left) ROC curve. (right) PR curve

### Prediction of gene expression change in the cell

There was the fifth Critical Assessment of Genome Interpretation (CAGI) competition at september 2019, and one of the challenges was to predict the effect of missense mutations on PTEN and TPMT protein stability ^41^. The protein stability change caused by mutations was measured by variant stability profiling (VSP) assay which detects the fluorescence from EGFP fused to the mutated protein. Therefore, the outcome of VSP indicates the gene expression of mutated proteins compared to wild-type proteins in the cell. According to the variant stability scores provided by CAGI, we assumed that variants with scores less than 0.5 are pathogenic while benign variants have scores higher than 0.5. As a consequence, there were 1,032 pathogenic variants and 2,848 benign variants for PTEN. In the case of TPMT, the number of pathogenic variants was 662 and that of benign variants was 3,094.

The result of performance using cross validation is described in Table 4 and Table 5. The result reassures the fact that multi-task learning with conservation data helps to make pathogenicity prediction to be accurate. Especially for these data, only if any types of knowledge transfer were applied, the performance of those models were generally superior to models using only one type of data. This may be because the pathogenic data used for training was relatively small and thereby conservation data was highly effective to reduce the overfitting. We also compared the performance with REVEL and PrimateAI for the same variants we tested for PTEN and TPMT (Figure 10). The superior performance of 3Cet indicates that transferring knowledge from evolutionary conservation is also effective to predict impacts of mutations in the cell.

**Table 4.**
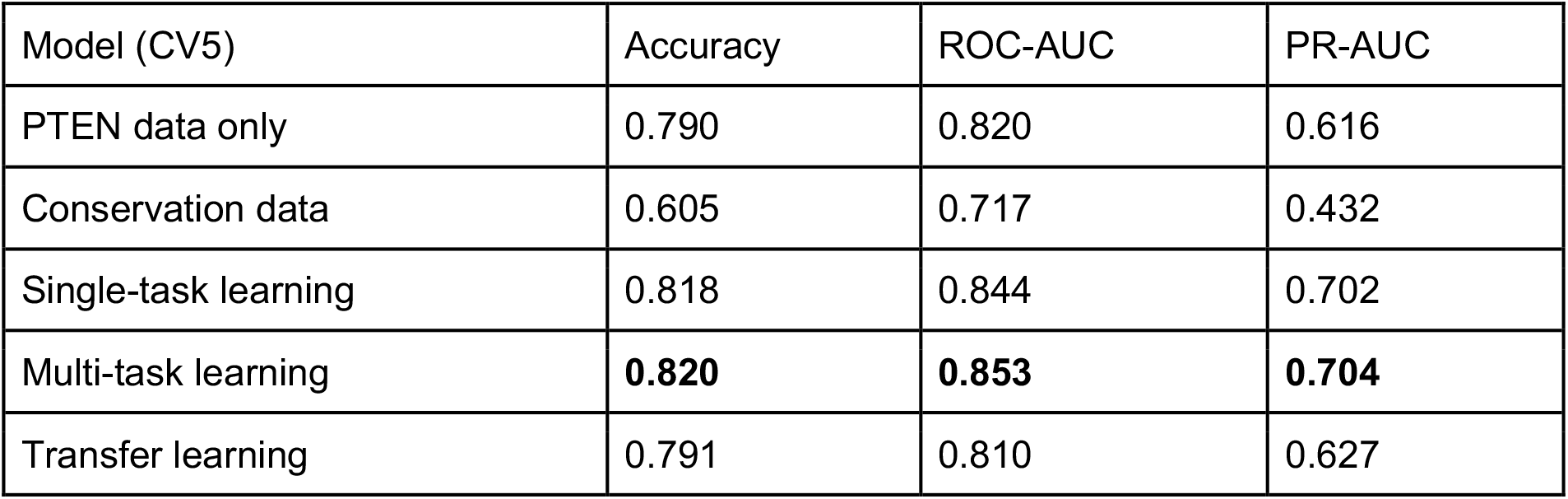
Accuracy, ROC-AUC, and PR-AUC of 5-fold cross validation for PTEN stability data

**Table 5.** Accuracy, ROC-AUC, and PR-AUC of 5-fold cross validation for TPMT stability data

| Model (CV5) | Accuracy | ROC-AUC | PR-AUC |
| --- | --- | --- | --- |
| TPMT data only | **0.814** | 0.729 | 0.369 |
| Conservation data | 0.732 | 0.728 | 0.388 |
| Single-task learning | 0.790 | 0.777 | 0.476 |
| Multi-task learning | 0.774 | **0.781** | **0.500** |
| Transfer learning | 0.784 | 0.773 | 0.488 |

**Figure 10.**
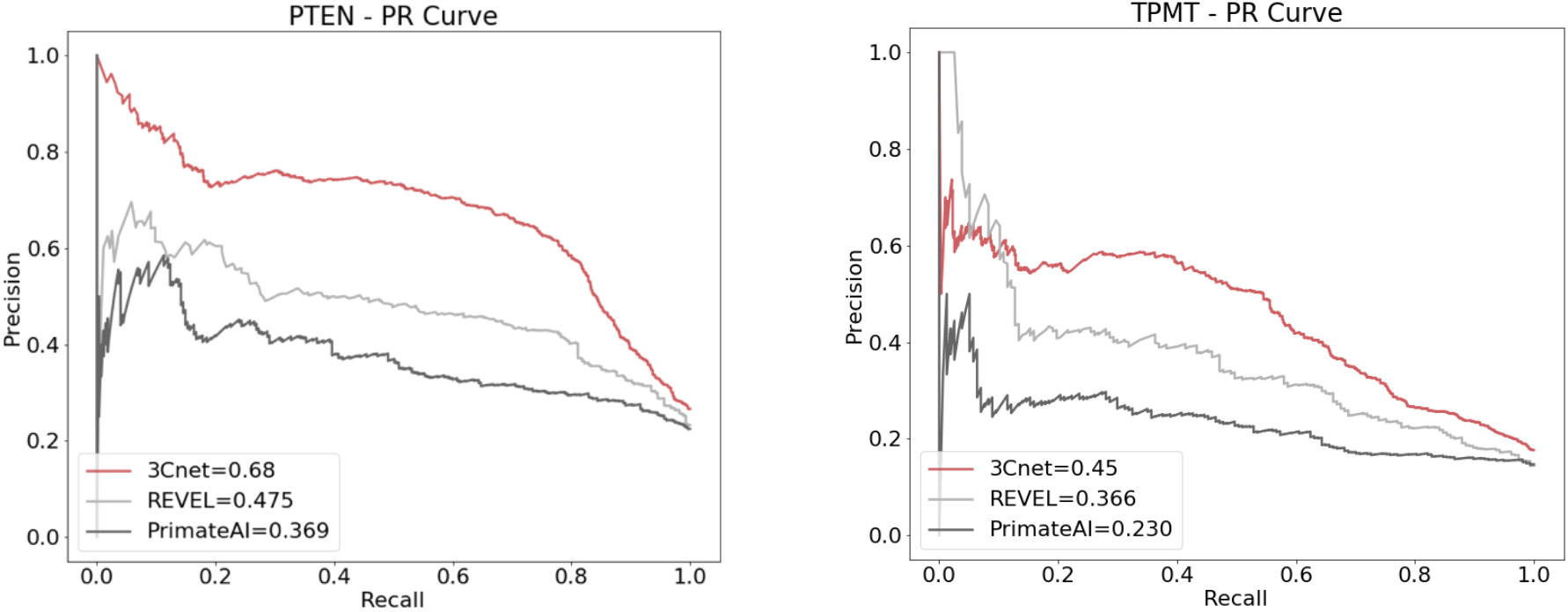
Precision-Recall curve for the prediction of gene expression changes in the cell. (left) PTEN. (right) TPMT

## Discussion

3Cnet, the first deep learning network using knowledge transfer to solve the pathogenicity prediction of human variants, has two strengths for clinical use. One is the accuracy of the model when classifying pathogenic variants and benign variants defined by the ACMG guideline. Variant classification using 3Cnet is nearly as accurate as ensemble based methods such as REVEL, even without making use of other scoring methods. The other is the capability of finding the disease-causing variant among a number of uncertain variants. 3Cnet found disease-causing variants of patients with higher sensitivity compared to other algorithms. Such a performance was possible because 3Cnet simultaneously trained various types of data, which were clinical data examined based on the ACMG guidelines, common variants found in the general population, and the conservation data generated based on the trinucleotide context and evolutionary conservation. Multi-task learning between clinical pathogenicity and evolutionary fatality, made it possible to train the pathogenic variants without overfitting despite the insufficient number of clinical data. Also, MSA features given as inputs of 3Cnet enabled the predictor to identify viable amino-acids for each residue conserved throughout the evolution. Moreover, we checked whether knowledge transfer also works for other types of pathogenicity prediction including protein stability changes and gene expression changes in the cell. The result is meaningful because it proves the fact that the conservation data can explain general impacts of mutations in the protein, in the cell, and in the human body. What makes it possible to train such sparse clinical data without biases was the general knowledge transferred from the conservation data.

Owing to the high performance of 3Cnet, the application of 3Cnet can be diverse depending on the purpose of researchers or medical doctors. One application would be finding the true disease-causing variant from the gnome of a patient. On average, there were around 100∼400 rare missense mutations within the genome of a patient ^42^. Among those mutations, only one or two mutations are known to cause the symptoms while the others are merely benign variants. Therefore, 3Cnet can be used to narrow down the candidate variants based on the scores thereby reducing time and cost spent for diagnosis. Thanks to the high recall rate, 3Cnet can help improving diagnostic rate significantly. However, relatively low precision for high thresholds needs to be enhanced for score-based diagnosis. One of the solutions would be to integrate known gene and protein information of which conventional prediction tools make use ^14^. In addition, scoring variants using 3Cnet can be used to find novel gene-disease association. Some genes may not be designated to any diseases in OMIM, a gene-disease mapping database ^43^, but they could have variants that seem to be pathogenic based on 3Cnet scores. If multiple pathogenic variants are found in a single gene and the symptoms of those variants are similar to a specific disease, we can estimate that the gene is correlated with the disease. Last but not least, the features extracted from the deep neural network can be utilized to train other deep learning models in which the pathological impact of a variant is important. For example, those features can be utilized to predict the functional domains of genes based on the sequence.

Even though 3Cnet can only predict the pathogenicity of missense variants at the moment, other variants such as nonsense variants or frameshift variants could be trained by 3Cnet. As 3Cnet makes use of the protein sequences as the input features and zero-padding can be applied for residues where the corresponding amino acids are missing, the network has the possibility to deal with different types of mutations other than missense mutations. Also, 3Cnet is the first RNN based model utilizing sequence inputs to predict pathogenicity. Instead of RNN, we can also make use of attention based networks such as a transformer to train the model. As a popular technique for natural language processing, transformers can train the context of a protein sequence and associate amino-acids at distant residues using the attention layer ^44^. Finally, as the input features of 3Cnet is the exon sequence, multiple nucleotide variants (MNVs) of a patient, with which the algorithms for single nucleotide mutations cannot deal, can be considered for patient diagnosis. In some cases, individual variants within MNVs seem benign but the integrated consequence is pathogenic, and vice versa. Such effects cannot be predicted using the pathogenicity prediction of single point mutations. Further data accumulation and extension of the network would make it possible to predict the integrated pathogenicity of multiple variants within a gene based on the overall sequence.

## References

1. Lek, M. et al. Analysis of protein-coding genetic variation in 60,706 humans. Nature 536, 285–291 (2016).

2. Homsy, J. et al. De novo mutations in congenital heart disease with neurodevelopmental and other congenital anomalies. Science 350, 1262–1266 (2015).

3. Iossifov, I. et al. The contribution of de novo coding mutations to autism spectrum disorder. Nature 515, 216–221 (2014).

4. Jin, S. C. et al. Contribution of rare inherited and de novo variants in 2,871 congenital heart disease probands. Nat. Genet. 49, 1593–1601 (2017).

5. Pérez-Palma, E., Gramm, M., Nürnberg, P., May, P. & Lal, D. Simple ClinVar: an interactive web server to explore and retrieve gene and disease variants aggregated in ClinVar database. Nucleic Acids Res. 47, W99–W105 (2019).

6. Gatz, C. et al. Identification of cellular pathogenicity markers for SIL1 mutations linked to marinesco-sjögren syndrome. Front. Neurol. 10, (2019).

7. Richards, S. et al. Standards and guidelines for the interpretation of sequence variants: A joint consensus recommendation of the American College of Medical Genetics and Genomics and the Association for Molecular Pathology. Genet. Med. 17, 405–424 (2015).

8. Landrum, M. J. et al. ClinVar: Public archive of interpretations of clinically relevant variants. Nucleic Acids Res. 44, D862–D868 (2016).

9. Liu, X., Jian, X. & Boerwinkle, E. dbNSFP: A lightweight database of human nonsynonymous SNPs and their functional predictions. Hum. Mutat. 32, 894–899 (2011).

10. Amendola, L. M. et al. Performance of ACMG-AMP Variant-Interpretation Guidelines among Nine Laboratories in the Clinical Sequencing Exploratory Research Consortium. Am. J. Hum. Genet. 98, 1067–1076 (2016).

11. Ghosh, R., Oak, N. & Plon, S. E. Evaluation of in silico algorithms for use with ACMG/AMP clinical variant interpretation guidelines. Genome Biol. 18, 1–12 (2017).

12. Rentzsch, P., Witten, D., Cooper, G. M., Shendure, J. & Kircher, M. CADD: predicting the deleteriousness of variants throughout the human genome. Nucleic Acids Res. 47, D886–D894 (2019).

13. Ph, R. O. Predicting the Functional, Molecular, and Phenotypic Consequences of Amino Acid Substitutions using Hidden Markov Models - supplement. 1–8.

14. Carter, H., Douville, C., Stenson, P. D., Cooper, D. N. & Karchin, R. Identifying Mendelian disease genes with the variant effect scoring tool. BMC Genomics 14 Suppl 3, S3 (2013).

15. Adzhubei, I. A. et al. A method and server for predicting damaging missense mutations. Nature methods vol. 7 248–249 (2010).

16. Kumar, P., Henikoff, S. & Ng, P. C. Predicting the effects of coding non-synonymous variants on protein function using the SIFT algorithm. Nat. Protoc. 4, 1073–1081 (2009).

17. Quang, D., Chen, Y. & Xie, X. DANN: a deep learning approach for annotating the pathogenicity of genetic variants. Bioinformatics 31, 761–763 (2015).

18. Grimm, D. G. et al. The evaluation of tools used to predict the impact of missense variants is hindered by two types of circularity. Hum. Mutat. 36, 513–523 (2015).

19. Sundaram, L. et al. Predicting the clinical impact of human mutation with deep neural networks performed the deep learning analysis. Nat. Genet. 50, 1161–1170 (2018).

20. Chen, T., Goodfellow, I. & Shlens, J. Net2Net: Accelerating learning via knowledge transfer. 4th Int. Conf. Learn. Represent. ICLR 2016 - Conf. Track Proc. 1–12 (2016).

21. Song, W. et al. Exploring the landscape of pathogenic genetic variation in the ExAC population database: insights of relevance to variant classification. Genet. Med. 18, 850– 854 (2016).

22. den Dunnen, J. T. et al. HGVS Recommendations for the Description of Sequence Variants: 2016 Update. Hum. Mutat. 37, 564–569 (2016).

23. Pruitt, K. D., Tatusova, T. & Maglott, D. R. NCBI Reference Sequence (RefSeq): a curated non-redundant sequence database of genomes, transcripts and proteins. Nucleic Acids Res. 33, D501–4 (2005).

24. Karczewski, K. J. et al. The ExAC browser: displaying reference data information from over 60 000 exomes. Nucleic Acids Res. 45, D840–D845 (2017).

25. Gilissen, C., Hoischen, A., Brunner, H. G. & Veltman, J. A. Disease gene identification strategies for exome sequencing. Eur. J. Hum. Genet. 20, 490–497 (2012).

26. Church, D. M. et al. Modernizing reference genome assemblies. PLoS Biol. 9, e1001091 (2011).

27. Remmert, M., Biegert, A., Hauser, A. & Söding, J. HHblits: Lightning-fast iterative protein sequence searching by HMM-HMM alignment. Nat. Methods 9, 173–175 (2012).

28. Suzek, B. E., Huang, H., McGarvey, P., Mazumder, R. & Wu, C. H. UniRef: comprehensive and non-redundant UniProt reference clusters. Bioinformatics 23, 1282– 1288 (2007).

29. Mikolov, T., Sutskever, I., Chen, K., Corrado, G. S. & Dean, J. Distributed Representations of Words and Phrases and their Compositionality. in Advances in Neural Information Processing Systems 26 (eds. Burges, C. J. C., Bottou, L., Welling, M., Ghahramani, Z. & Weinberger, K. Q.) 3111–3119 (Curran Associates, Inc., 2013).

30. Hochreiter, S. & Schmidhuber, J. Long Short-Term Memory. Neural Comput. 9, 1735– 1780 (1997).

31. Ruder, S. An Overview of Multi-Task Learning in Deep Neural Networks. (2017).

32. Eraslan, G., Avsec, ž., Gagneur, J. & Theis, F. J. Deep learning: new computational modelling techniques for genomics. Nat. Rev. Genet. 20, 389–403 (2019).

33. Taylor, L. & Nitschke, G. Improving Deep Learning using Generic Data Augmentation. (2017).

34. Bleeker, S. E. et al. External validation is necessary in prediction research: a clinical example. J. Clin. Epidemiol. 56, 826–832 (2003).

35. Stenson, P. D. et al. Human Gene Mutation Database (HGMD): 2003 update. Hum. Mutat. 21, 577–581 (2003).

36. Seo, G. H. et al. Diagnostic yield and clinical utility of whole exome sequencing using an automated variant prioritization system, EVIDENCE. Clin. Genet. (2020) doi:10.1111/cge.13848.

37. Ozenne, B., Subtil, F. & Maucort-Boulch, D. The precision--recall curve overcame the optimism of the receiver operating characteristic curve in rare diseases. J. Clin. Epidemiol. 68, 855–859 (2015).

38. Sasidharan Nair, P. & Vihinen, M. VariBench: A Benchmark Database for Variations. Hum. Mutat. 34, 42–49 (2013).

39. Chen, C.-W., Lin, J. & Chu, Y.-W. iStable: off-the-shelf predictor integration for predicting protein stability changes. BMC Bioinformatics 14 Suppl 2, S5 (2013).

40. Montanucci, L., Martelli, P. L., Ben-Tal, N. & Fariselli, P. A natural upper bound to the accuracy of predicting protein stability changes upon mutations. Bioinformatics 35, 1513–1517 (2019).

41. Pejaver, V. et al. Assessment of methods for predicting the effects of PTEN and TPMT protein variants. Hum. Mutat. 40, 1495–1506 (2019).

42. Auton, A. et al. A global reference for human genetic variation. Nature 526, 68–74 (2015).

43. Amberger, J. S. & Hamosh, A. Searching Online Mendelian Inheritance in Man (OMIM): A Knowledgebase of Human Genes and Genetic Phenotypes. Curr. Protoc. Bioinforma. 58, 1.2.1-1.2.12 (2017).

44. Vaswani, A. et al. Attention is all you need. Adv. Neural Inf. Process. Syst. 2017-December, 5999–6009 (2017).

